# Coupling differential adhesion to cell signaling avoids kinetic traps to yield robust multicellular self-organization

**DOI:** 10.1101/2025.10.23.684104

**Authors:** Guy Pelc, Wesley L. McKeithan, Yipei Guo, Michael P. Brenner, Wendell A. Lim, Mor Nitzan

**Affiliations:** School of Computer Science and Engineering, The Hebrew University of Jerusalem, Jerusalem, Israel; Cell Design Institute, Dept. of Cellular and Molecular Pharmacology, University of California, San Francisco, CA; School of Engineering and Applied Sciences, Harvard University, Cambridge; Program in Biophysics, Harvard University, Cambridge; Institute of High Performance Computing, Agency for Science, Technology and Research, Singapore; Physics Department, Harvard University, Cambridge; Racah Institute of Physics, The Hebrew University of Jerusalem, Jerusalem, Israel; Faculty of Medicine, The Hebrew University of Jerusalem, Jerusalem, Israel

## Abstract

Differential adhesion, where cells physically reorganize based on their heterogeneous adhesion preferences, is one of the major models for self-organization in development and tissue formation. However, accumulating evidence suggests that differential adhesion is many times insufficient for robust convergence to a target minimal energy multicellular structure. Here we use computational simulations and engineered synthetic cell circuits to systematically explore alternative mechanisms for programming formation of a simple two-cell type core-shell morphology. Starting with two pre-differentiated cell types with constitutively high differential adhesion leads to kinetic trapping in variable, multi-core structures. In contrast, hybrid mechanisms that gradually induce differential adhesion upon cell-cell contact signaling consistently converge to the target single-core structure, in a manner robust to variation in cell numbers, interaction energy, and noise. This work delineates intrinsic limitations of self-organizing systems based solely on differential adhesion, and shows how inducible systems provide a way to invoke the strong adhesion required to maintain a multicellular structure, while avoiding the pitfall of kinetic traps. This study illustrates how joint computational and experimental exploration of synthetic circuits can be used to probe key developmental principles and tradeoffs and inform the design of synthetic development and self-organization.

## 1 Introduction

One of the major conceptual paradigms thought to drive self-organization in organismal and tissue development is physical cell sorting based on differential adhesion, where cell types exhibiting heterogeneous adhesion preferences spatially reorganize to minimize their interfacial free energy [1, 2, 3]. Differential adhesion has been suggested to play a key role in the formation of tissue structure in diverse biological contexts, including tissue patterning in the Drosophila retina [4], the mammalian olfactory epithelium [5, 6] and mammalian auditory epithelium [7], as well as the formation of stable boundaries between cell types and tissues in the developing vertebrate hindbrain [7, 8] and limb regeneration [9]. However, pattern formation driven by differential adhesion, and specifically cadherin-dependent sorting, can be limited and can yield variable, local minima configurations under certain parameter regimes, rather than converging on a single minimal energy configuration. For example, in engineered populations of mammalian cells of two differentially-adhesive cell types, while small (1,000 cells) 3D multicellular systems converge to stable two-domain configurations, large (10,000 cells) systems get dynamically trapped in variable multi-domain configurations [10, 11]. Several recent works have shown evidence that regulated adhesion may be crucial for robust and convergent self-organization. For example, in the context of early embryonic development in mice, where epiblast mesendoderm progenitors segregate to mesoderm and definitive endoderm (DE), it was found in vitro that the expression of the cell adhesion molecules E-cadherin increases simultaneously with physical sorting of cells expressing DE regulators (Sox17+ cells), where the aggregation of Sox17+ cells resulted in the upregulation of E-cadherin, yet the mechanism for its delayed upregulation is still not fully understood [12]. This finding is consistent with observations in the embryo showing that high levels of E-cadherin were found in DE cells only after they had intercalated into the visceral endoderm [13, 14]. Therefore, it was suggested that the gradual induction of E-cadherin may prevent premature formation of multiple DE cell clusters within the epiblast [12].

The self-assembly of abiotic interacting molecules or particles (e.g. DNA origami, DNA-coated colloidal particles and magnetic handshake materials) has been systematically explored experimentally to understand fundamental properties such as efficiency, scalability, and limits of control of size and shape of self-assembled structures [15, 16, 17]. However, these abiotic systems differ from biological self-organizing systems, such as those observed in development, which can, in principle, harness regulated changes in physical interactions. This gap is bridged by recent advancements in synthetic biology that have opened new possibilities for programming cells to form complex multicellular structures by engineering cell identities, capabilities, and signaling pathways [18, 11, 19, 20, 21, 22]. In particular, recent synthetic biology studies have established that cell signaling can be programmed to alter the properties of physical interactions and thus alter the formation of multicellular configurations [23, 22].

Here we use computation and synthetic biology to demonstrate the limitations of multicellular self-organizing systems that are based on adhesion alone and their dynamic divergence as they tend to become trapped in local configuration minima. We show that gradual increase in adhesion can be a general solution to more robust self-organization, and that cell-cell signaling which induces adhesion is a possible mechanism underlying such solution, which is both experimentally and evolutionarily accessible. We show how adding cell-cell signaling to generate inducible adhesion enhances the robust physical organization of cells, and discuss how the induction mechanism can be used to control kinetics and converge to a core-shell structure in the context of synthetic development experiments, even in regimes where the dynamics of physical sorting by differential adhesion alone would likely yield heterogeneous, multiple-core configurations.

## 2 Results

### 2.1 Computational self-organization of core-shell multicellular configurations by differential adhesion leads to kinetic trapping in variable structures

We focus on ways to specify formation of one of the simplest possible multicellular target structures, a core-shell configuration, with one cell type forming a central core, and a second cell type forming a surrounding spherical shell (Figure 1A).

**Figure 1:**
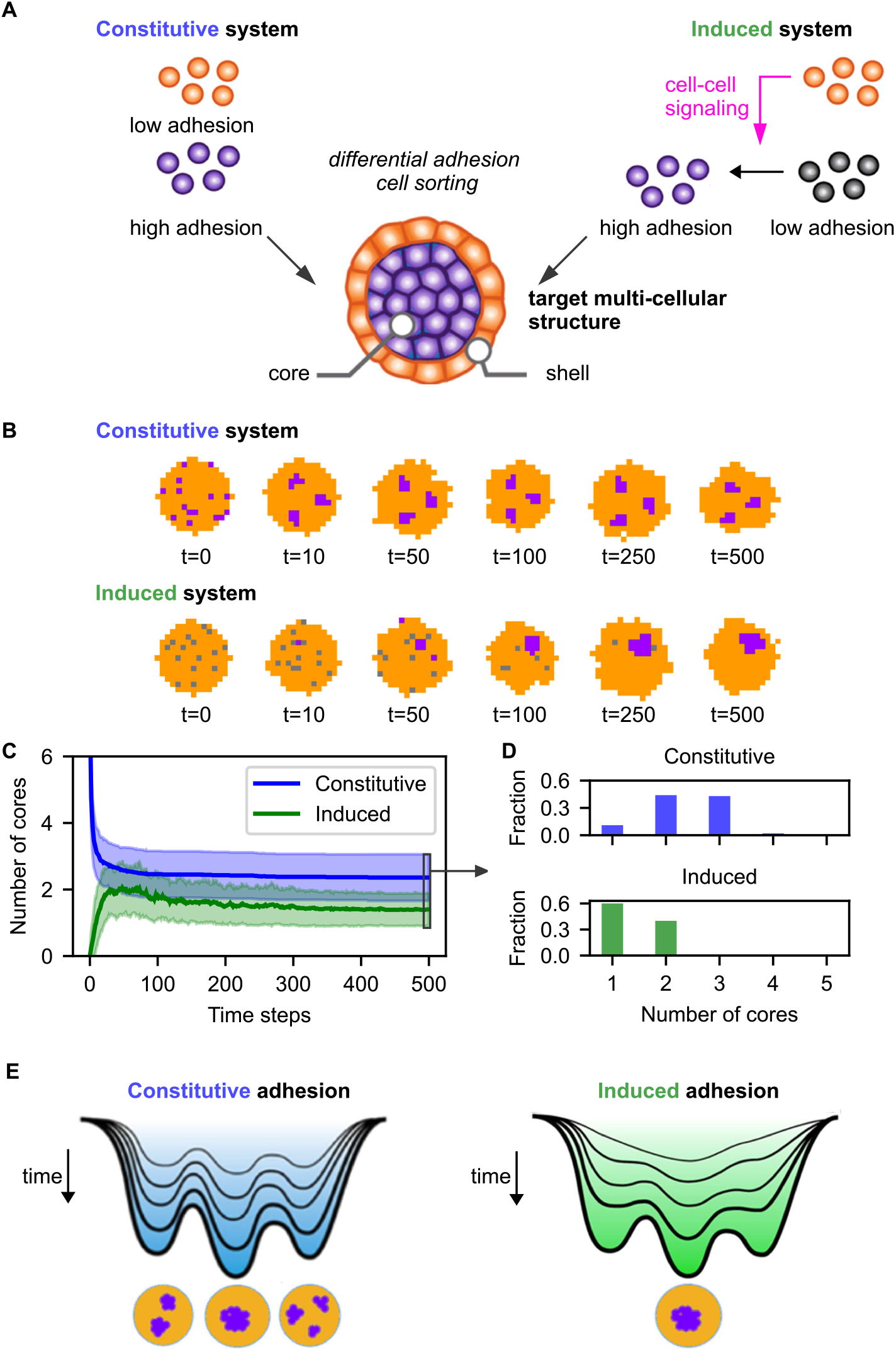
Modeling predicts that induced differentiation coupled to physical sorting yields more robust formation of single core-shell morphologies, while constitutive adhesion yields variable multi-core morphologies. **(A)** Schematic of the constitutive vs. induced circuits. **(B)** Snapshots of representative simulated temporal trajectories of the two circuits in the Potts model. Type-A cells are shown in orange, low-adhesion B cells in gray, and high-adhesion B cells in purple. **(C)** Number of cores over time for the two circuits (lines: mean, shaded area: std). **(D)** Histogram of the number of cores at the final time step for the two circuits show that the induced system tends to converge onto fewer cores. **(E)** Schematics of the energy landscapes corresponding to the multicellular configurations of the constitutive and induced adhesion systems, summarizing a model for how the induced system may favor more robust convergence to single-core structures, while the constitutive system can lead to kinetic trapping in multi-core structures. While the energy landscape of the constitutive system remains constant throughout the self-organization process (progression of time is represented on the vertical axis), for the induced system there is a gradual transformation from a relatively flat to a rugged energy landscape, corresponding to the gradual induction of adhesive cells. Results are shown for 100 instantiations, number of cells = 150, fraction of core cells = 0.1, *T* = 5, binding energy: A-A, A-B = 10, B-B = 100, *p* = 0.01, and Monte Carlo simulation time steps = 500. See Section 4.1.1 for implementation details.

Core-shell configurations emerge in diverse developmental and homeostatic contexts, including Hydra regeneration [24], avian gastrulation [25], and core-mantle morphology of endocrine cells in pancreatic islets [26]. First principles dictate that a core-shell structure is the minimum energy structure for a system of two cell types (A,B) with differential self-adhesion: the cell type (B) with stronger self-adhesion sorts to the core, while the cell type with weaker self-adhesion (A) sorts to the shell.

To examine this, we construct a computational model, combining aspects from cellular automata and cellular Potts model (Methods 4.1.1, Figure 1A,B). We first considered a constitutive system where two cell types (A and B) have fixed, predetermined adhesion properties: B-type cells adhere much more strongly to each other than to A-type cells, or than A-type cells adhere to themselves (Figure 1A, left). We initialize the system as a spheroid with a random mixture of A and B cells. While the lowestenergy, stable configuration of this system has a core of B cells surrounded by a shell of A cells, in 89 of the 100 realizations of our simulations the cells self-organize into a higher energy structure with multiple cores of B cells (median number of cores: 2; mean: 2.36, std: 0.70), surrounded by A cells (Figure 1C,D).

These configurations are local minima of the energy landscape associated with the model, as opposed to the global minimum configuration of a single core (Figure 1E).

### 2.2 Induced differentiation by cell-cell signaling promotes robust computational self-organization

As an alternative mechanism, we considered a system in which differential adhesion of inactivated core cells was gradually induced (Methods 4.1.1, Figure 1A,B). Such an induced system exhibited more stable convergence to single-core structures (60/100 realizations) relative to the constitutive system (11/100 realizations, Figure 1C,D). One accessible biological mechanism that can generate gradual adhesion induction is based on cell-cell contact signaling, and therefore, we considered an induced system where cellular sorting is intrinsically coupled to induced differentiation. In this case, A and B cells start out both having equal low adhesive properties (neutral). However, B cells are activated to become highly self-adhesive B cells (identical to the B cells in the constitutive system) upon local interaction with A cells, via A→B cell-cell signaling (Figure 1A, right). In this induced model, we observe far more robust self-organization: while a typical temporal trajectory of the constitutive system yields a multiple core configuration (median number of cores: 2; mean: 2.36, std: 0.70), the induced system tends to converge to a single core configuration of activated B cells (median number of cores: 1; mean: 1.4, std 0.49), and is associated with lower-energy configurations of the model (Supplementary Figure S1A).

Our findings, which were robust across a wide range of effective temperatures (representing noise, Supplementary Figure S1B), demonstrate that gradual activation of B-type cells in the induced system enables robust convergence to the core-shell global minimum configuration, whereas the constitutive system often gets trapped in local minima with multiple cores (Figure 1E).

### 2.3 Stochastic cluster model

While the Potts model offers detailed single-cell resolution, it does not support the drifting of cell clusters as a single entity. Additionally, its high resolution comes with a computational cost, which limits the number of cells that can be modeled. To address these limitations, we introduced a stochastic cluster model to generalize system dynamics and characterize, at larger-scales and extended dynamical regimes, the distinction between the constitutive and induced systems (Figure 2A). In this abstraction, high-adhesion cell clusters move within a background of low-adhesion cells, and merge upon contact. This approach prioritizes essential dynamics by focusing on cluster-level observables, significantly reducing computation time (Figure 2A, Table 1, Methods 4.1.2).

**Table 1.** Comparison of the Potts model and the stochastic cluster model (Figure 2A)

| Potts model | Stochastic cluster model |
| --- | --- |
| Single-cell resolution | Cluster-level resolution |
| Models all cell types | Explicitly models high-adhesion cells only |
| Movement via position swapping | Movement as random walks |
| Speed influenced by temperature and interaction strength | Speed decays as a power law with respect to the cluster’s radius, controlled by a varying $x$ parameter: $s \propto r^{-x}$. The speed decay factor $x$ represents the system’s implicit dependence on temperature and binding energies. |
| High adhesion activation through A→B cell-cell signaling | High adhesion stochastic activation at interval $k$ |

**Figure 2:**
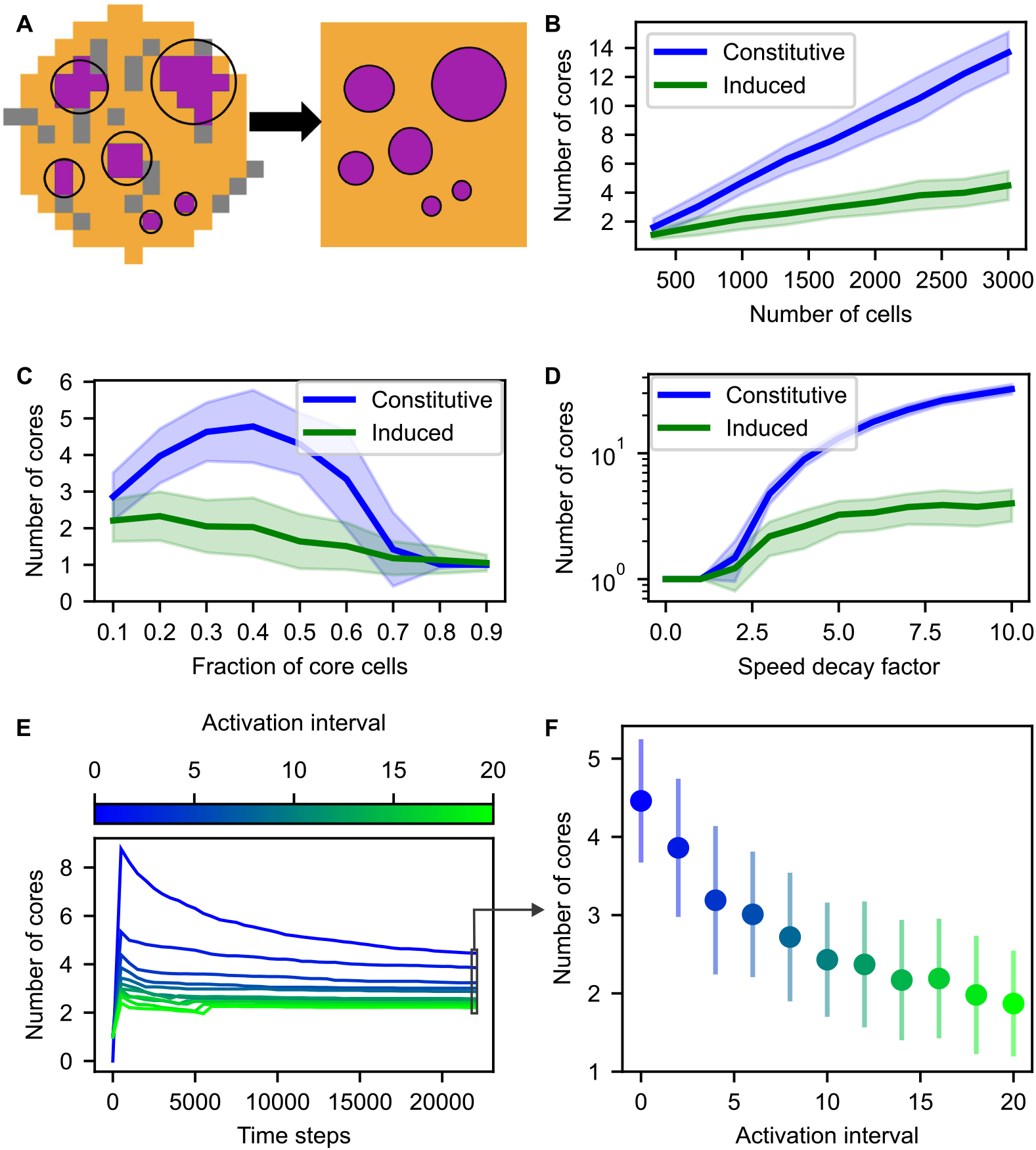
Simulations with the stochastic cluster model reveal the robustness of the induced system compared to the constitutive system across a broad parameter range. **(A)** Schematic comparison of the Potts model and the stochastic cluster model, where low-adhesion A (B) cells are in orange (gray), and high-adhesion B cells are in purple. See Table 1 for details. **(B**,**C)** Number of cores in the constitutive (blue) and induced (green) systems as a function of the total number of cells (B) and the fraction of core cells (C) (lines: mean, shaded area: std). **(D)** The number of cores increases with the speed decay factor *x*, where the speed *s* of a cluster as a function of its radius *r* is modeled as *s ∝ r*^−*x*^. The induced system demonstrates a growing advantage over the constitutive system as *x* increases, which can be interpreted as the emergence of kinetic traps. **(E)** Evolution of the number of cores over time for different activation intervals *k*, representing the gradual nature of the activation process for high-adhesion cells in induced systems. **(F)** Final number of cores as a function of the activation interval *k*. Increasing the activation interval *k* between high-adhesion cell activations improves convergence to fewer cores, generalizing the constitutive system (*k* = 0) and induced system (*k >* 0). Default parameters: total number of cells = 1000, fraction of core cells = 0.3, *k* = 15, *x* = 3, number of instantiations = 100. See Section 4.1.2 for implementation details.

Each cluster performs a random walk with a speed that decreases as the cluster size increases, following a power-law decay (Methods 4.1.2). This relationship was derived from high-resolution, detailed simulations conducted using the Morpheus computational framework [27] (Supplementary Figure S7). In the induced model, B-type cells stochastically emerge from the background at a controlled time interval *k* (Methods 4.1.2).

The stochastic cluster model demonstrates clear computational advantages, allowing effective simulation of thousands of cells. For example, we measured a 10-fold decrease in runtime for the stochastic cluster model compared to the Potts model using default parameters and 3,000 cells.

Figure 2B illustrates the induced system’s superiority in achieving fewer cores, especially as the cell count increases. When considering varying fractions of core cells (Figure: 2C) the performance gap is more nuanced: systems with very low or very high core cell fractions exhibit diminished differences between induced and constitutive dynamics, as there are fewer kinetic traps in these regimes. More importantly, the stochastic cluster model reveals a tradeoff between the systems. The decay factor *x*, which determines how cluster speed decreases as a function of its radius (*s ∝ r*^−*x*^), plays a critical role. For small decay factors (*x <* 1), both systems converge to single cores, though the induced system achieves this slower (Figure S2) due to the time it takes for all B cells to become activated. As *x* increases, the constitutive system stabilizes at significantly higher core counts compared to the induced system (Figure 2D).

This can be interpreted as the emergence of kinetic traps with increasing *x*, where clusters form in multiple regions and cannot merge within the simulation time due to their decreased speed. In the induced system, this effect is much less pronounced, as single cluster cells that appear gradually are not affected by the decay and can potentially merge with the main core before forming a separate cluster with another single cell.

Finally, by varying the activation interval *k*, we transition smoothly from induced to constitutive dynamics. Lower *k* values result in less gradual activation, reducing to constitutive behavior at *k* = 0. Higher *k* values enhance convergence to single-core configurations, emphasizing the critical role of gradual activation (Figures 2E,F).

### 2.4 Engineered interacting cells exhibit robust self-organization with signaling-induced differential adhesion relative to constitutive adhesion

To experimentally test the hypothesis that induced differential adhesion is more robust, we use synthetic biology to systematically engineer cells with constitutive and induced adhesion (Figure 3A). We engineered mouse L929 fibroblasts which were placed in low adhesion U-bottom wells, and followed their dynamic self-organization over 21 hours at 1-hour intervals with automated confocal microscopy. The imaging output then generated 3D reconstructions of the multicellular structure (Figure 3B). Since L929 cells do not naturally self-organize, but form a random aggregate, any particular spatial organization could be attributed to the engineered synthetic circuits. For both the induced and constitutive cases, we designed experiments for different initial numbers of cells (300 and 900 cells), different A-to-B cell type ratios (1:9, 3:7, 1:1, 7:3, 9:1), and 4 replicates for each experiment (Methods 4.2, 4.3).

**Figure 3:**
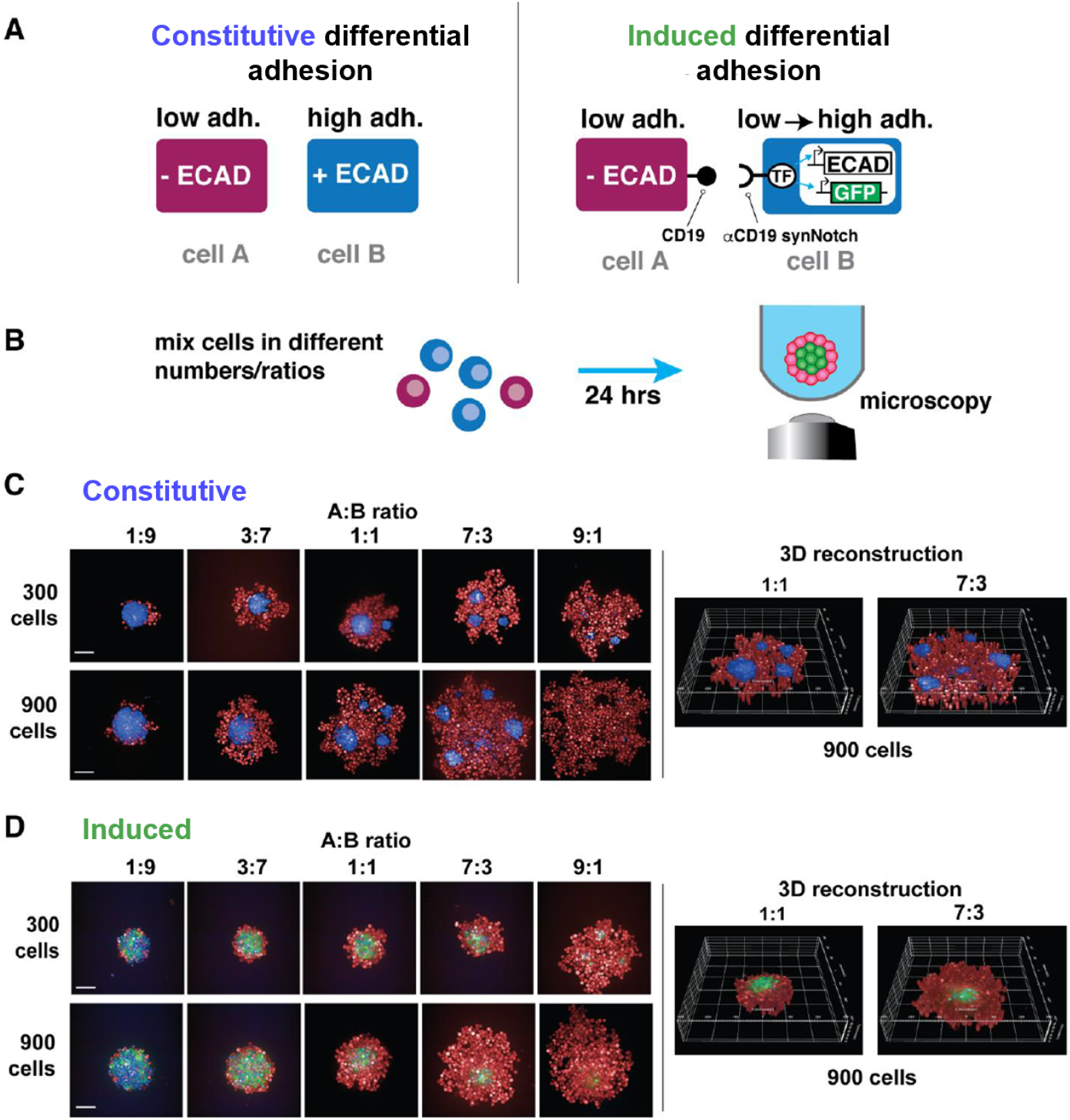
Experimental studies with synthetic core-shell systems show more stable convergence to a single core configuration of induced-adhesion compared to constitutive systems. **(A)** Design of constitutive vs induced synthetic differential adhesion circuits. For the constitutive system, low adhesion cells (labelled with IFP) are mixed with high adhesion cells that constitutively express Ecad (labelled with BFP). For the induced system, sender cells (labelled with BFP) express the synNotch ligand CD19, and receiver cells (labelled with IFP) express the cognate anti-CD19 synNotch receptor along with its response element. Ecadherin (Ecad) and GFP genes are placed under the control of the synNotch responsive promoter in the receiver cells. **(B)** Experimental design - cells are mixed in low adhesion, U bottom wells, and observed by microscopy for 21 hours. Final time experimental snapshot as well as 3D reconstructions for Constitutive **(C)** and Induced **(D)** circuits, with different initial cell numbers (300 and 900 cells), and different cell ratios (non-core to core ratio: 1:9, 3:7, 1:1, 7:3, 9:1). Images were false colored in this figure for consistency so that all low adhesion cells appear red.

For the constitutive system, we mixed low adhesion cells (A-type cells; labelled with IR fluorescent protein, IFP) with high adhesion cells that constitutively express Ecad (B-type cells; labelled with blue fluorescent protein, BFP) (Figure 3A, left). In this case, the B-type cells sort to form a single or multiple core structures surrounded by A-type cells (Figure 3C). For the induced system, we used the modular synthetic notch (synNotch) juxtacrine signaling system to engineer genetic programs in which specific cell-to-cell contacts induce increase in cadherin-dependent cell adhesion [28, 23]. We engineered a sender cell (A-type cells; labelled with BFP) expressing the synNotch ligand CD19, and a receiver cell (B-type cells; labelled with IFP) expressing the cognate anti-CD19 synNotch receptor along with its response element. We placed Ecadherin (Ecad) and green fluorescent protein (GFP) genes under the control of the synNotch responsive promoter in the receiver cells (Figure 3A, right). As we previously showed [23], when mixing A-type and B-type cells, B-type cells express GFP and Ecad, which leads to their sorting towards the middle of the spheroid. This self-organization process eventually leads in the majority of cases to a 2-layered spheroid, consisting of a core of B-type cells surrounded by a shell of A-type cells (Figure 3D).

We find that after 21 hours, consistent with our computational predictions, the system where differentiation is induced by cell-cell signaling forms a two-layered core-shell structure with a single core across a broad range of experimental scenarios involving variable cell numbers and cell-type ratios (Figure 3D). In contrast, the constitutive system tends to form a spheroid of A-type cells containing multiple cores of B-type cells (Figure 3C).

Snapshots of the time dynamics reveal distinct behaviors between the two systems. The constitutive system rapidly yields multiple cores (Figure 4A), whereas the induced system gradually converges to a single core (Figure 4B). Systematic analysis of the dynamics of the number of cores (Figure 4B,C, Methods 4.4) confirms this trend, with substantially lower variance in the number of cores for the induced system across all experimental conditions and trials (Figure 4B,C, Supplementary Figure S5).

**Figure 4:**
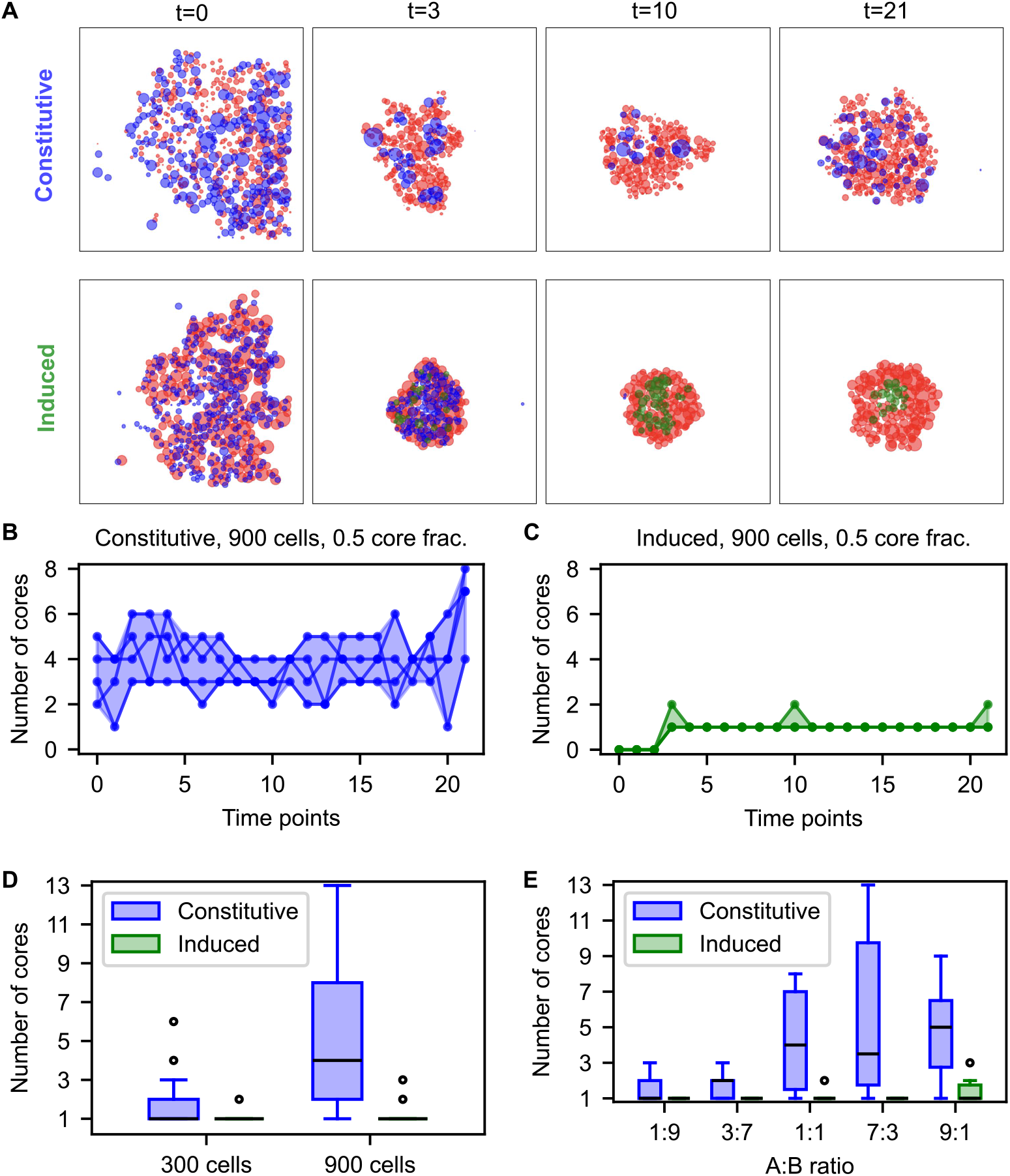
Experimental comparison of synthetically engineered constitutive vs induced adhesion self-organized circuits confirms that induced structure formation is more robust, and converges within several hours to a core-shell structure. **(A)** experimental snapshots of the constitutive and induced self-organization over time with 900 cells and 0.5 fraction of core cells (constitutive: low adhesion A cells in red, high adhesion B cells in blue; induced: low adhesion A/B cells in red/blue, high adhesion B cells in green). **(B**,**C)** Number of cores over time with 900 cells and 0.5 fraction of initial core cells, for constitutive (B) and induced (C) systems (lines: single trial trajectories; shaded area: range of values at a given timepoint). **(D**,**E)** The number of cores in the constitutive (blue) and induced (green) systems as a function of the initial total cell number (D), and as a function of the fraction of core cells (E). The boxes extend from the lower to upper quartile values of the data, with a line at the median, whiskers extend from *Q*3 + 1.5 · (*Q*3 − *Q*1) to *Q*1 − 1.5 · (*Q*3 − *Q*1), where *Q*1 and *Q*3 are the first and third quartiles, and circles represent outlier points.

Quantitatively, the induced system stabilizes its multicellular structure after approximately 5.8 *±* 3.3 hours, compared to 1.9 *±* 0.8 hours for the constitutive system (Supplementary Figure S5). Upon stabilization, the induced system achieves a significantly lower average distance from the center of mass (67.4 *±* 31.7 µm) compared to the constitutive system (108.8 *±* 38.4 µm) (Supplementary Figure S6).

Core counting statistics after 21 hours further highlights the differences between the systems. The induced system consistently forms fewer cores (median: 1; mean: 1.45; std: 0.97), with 75% of experiments resulting in a single core (30/40 trials), compared to the constitutive system, which forms more cores (median: 3; mean: 5.8; std: 4.85), with only 15% of trials leading to a single core (6/40 trials). This difference becomes particularly pronounced at higher cell numbers and lower core cell fractions (Figure 4D,E).

In the constitutive system, the number of cores increases with total cell numbers (Figure 4D) and decreases with the fraction of core cells (Figure 4E), aligning with computational predictions when core cell numbers are sufficient to form multiple cores (Figure 2B,C). In contrast, the induced system demonstrates robustness, consistently forming fewer cores than the constitutive system across all tested cell numbers and A-B cell-type ratios, emphasizing its stability and reliability.

These findings are also consistent with recent results by Tordoff et al. [29] who showed, computationally and experimentally, that the number of cores in their analogous constitutive system (composed of CHO K1 cells expressing low levels of cadherin, and HEK293FT cells expressing E-cadherin and N-cadherin and are more self-adhesive) increases with decreasing fraction of adhesive cells and with increasing total cell number.

## 3 Discussion

Coupling of differential adhesion-based sorting with cell-cell signaling, which induces differentiation, generates a continuous feedback in which cell identity regulates its location which in turn induces local interactions that regulate its identity. Here we provide a combined computational and experimental approach that suggests that such coupling between sorting and induced differentiation in a basic multicellular building block of development stabilizes self-organization, and leads to a more robust and reproducible convergence into a 3D core-shell configuration. We found induced differentiation to be a mechanism that can facilitate increased robustness of self-organization that avoids kinetic traps associated with constitutive differential adhesion. Taking a synthetic development perspective, changing parameters such as the adhesion strength, the effective temperature or cellular motility levels for cell sorting are generally challenging to control/program directly in experiments. In this work, we suggest an additional knob we can turn for synthetic development; adding the induction mechanism extends the experimental regime in which structural multicellular convergence is stable. Specifically in our setting, we extend the regime in which the constitutive, differential adhesion driven system converges to a core-shell structure by the complementary cell signaling induction mechanism.

In general, with increasingly complex body and tissue plans, we expect that higher numbers of cell types and heterogeneous binding profiles will lead the energy landscape to become more rugged, with more potential alternative stable spatial configurations. While such kinetic traps can be exploited in some systems to stabilize patterns that are not global energy minima such as periodic patterning (e.g. spots), here we focus on the case where the global minimum is the desired configuration. The induction mechanism facilitates a gradual transformation of the energy landscape, and can drive a developing biological system to robustly converge to a unique 3D configuration (Figure 1E). As organisms develop in complex and noisy environments, this kind of robust convergence is likely to be critical. This work clearly delineates the intrinsic limitations of self-organizing systems based solely on differential adhesion: the strength of constitutive adhesion interactions makes such systems prone to becoming trapped in local minima, a pitfall that is exacerbated with more cells and cell types. Inducible adhesion provides a way to invoke the strong adhesion required to maintain a multicellular structure, while avoiding kinetic traps. While there may exist multiple potential solutions to generate inducible adhesion, the one that is activated by cell-cell interactions is simple and accessible both experimentally, as we show in this study, and evolutionarily, which may explain why many developmental systems integrate cell-cell signaling regulation with changes in adhesion. This solution merges two of the main mechanisms for selforganization in development: differential adhesion-based physical sorting (where cell identity determines their position in the final stable assembly) and signaling-induced differentiation (whereby paracrine or juxtacrine signals from other cells provide positional information specifying resulting cell types, namely, the position of cells determines their identity [30, 31, 32, 33]).

While our results demonstrate the advantages of coupling differential adhesion to cell-cell signaling, this study has limitations. First, it focuses on the core-shell target structure, which consists of two cell types in a spherically symmetric configuration, representing a simplified scenario. Second, the experimental system relies on synthetic circuits that are isolated from external influences, such as external mechanical constraints, extracellular gradients, or interactions with surrounding tissues. Future work could explore how a gradual increase in complexity improves convergence in more complex target synthetic structures, such as the circuits introduced by Toda et al. [23], or in controlling organoid development [34]. Additionally, examining natural biological systems with varying adhesion and signaling mechanisms could offer valuable insights into whether gradual differential adhesion mechanisms play a role in developmental processes.

We believe that the induced adhesion mechanism is a special case of gradual lock-in biological processes that aid complex systems to converge onto desired spatial states. The combination of regulated differentiation and adhesion-based sorting, which is more feasible in biotic vs abiotic self-organizing systems, provides a route for dynamically generating more robust and complex structures. More generally, we show how computational and synthetic biology approaches can be combined to systematically probe specific design principles of development.

## 4 Methods

### 4.1 Computational multicellular self-organization model

#### 4.1.1 Cellular Potts model

There are two main aspects of our computational model for multicellular self-organization: physical sorting of the cells and differentiation (or state changes), both dependent on the immediate cellular environment of the cells. The first model we propose combines elements from a cellular automaton for the state changes and a cellular Potts model for the movement of cells (a.k.a Glazier-Graner-Hogeweg model) [35, 36], which was shown to faithfully capture biological pattern formation processes including cellular sorting (e.g. [37, 38, 39]).

In our first model, *N* cells are initiated on a 2D grid of size *L× L* within a circle. Each cell *i* is assigned at random a type *σ*(*i*), where *σ ∈* 1, 2, …, *m. m* and the relative fractions of cell types is set to correspond to the experimental setting we aim to model. As the simulation evolves in time, the cells can move to adjacent empty grid vertices, exchange places with their neighboring cells, and change their type in response to their cellular neighbors, or microenvironment. Specifically, starting from a certain initial configuration, at each step we choose at random a single grid site (uniformly at random over all occupied grid sites) and follow two steps:

1. State transition. If a cell at site *i* is an unactivated, low-adhesion B-type cell, it transitions to an activated, high-adhesion B-type cell with a certain probability *p*, provided that at least three of its neighbors are of type A.
2. Cellular movement using Metropolis-style updates. The cell at site *i* (of type *σ*(*i*)) switches places with the cell at site *j*, where *j* is chosen uniformly at random from nearest neighbors of *i* on the grid that are of a different type *σ*(*i*) *≠ σ*(*j*). The switch occurs with the following probability:

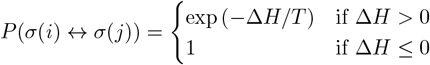

The energy *H* of a certain multicellular configuration is

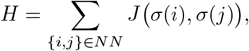

where *J* (*σ*(*i*), *σ*(*j*)) is the adhesion energy per unit contact area between cell types *σ*(*i*) and *σ*(*j*). Δ*H* is the energy difference associated with the potential cellular switch, and *T*, the effective temperature, is a positive constant.

Default simulation values: number of cells = 150, fraction of core cells = 0.1, *T* = 5, binding energy: *A* −*A, A*− *B* = 10, *B* −*B* = 100. *p* = 0.01.

Simulation time steps: = 500 Monte Carlo steps (MCS), where one MCS is equivalent to *N* simulation steps.

#### 4.1.2 Stochastic cluster model

The stochastic cluster model captures the dynamics simulated by the cell sorting module of the Morpheus modeling environment [27]. Morpheus employs an advanced version of the Potts model at sub-cellular resolution, incorporating additional parameters such as cell volume and surface constraints. These features enable highly detailed simulations that closely replicate biological cell movement. However, this level of detail comes at the cost of high computational demands, restricting the number of cells that can be simulated simultaneously.

By running multiple Morpheus simulations and tracking the movement of clusters of varying sizes, we observed that their center-of-mass trajectories follow a random walk. Moreover, the average speed *s* of these clusters obeys a power law, *s ∝ r*^−*x*^, where *r* is the effective cluster radius (Supplementary Figure S7), and the decay factor *x* depends on temperature and binding energy conditions.

In our stochastic cluster model, we simplify the system simulated by the Morpheus framework by omitting sub-cellular resolution and detailed parameters. This abstraction enables direct control over effective parameters and system observables, significantly reducing computational costs and facilitating exploration of a broader phase space, including large variations in cell numbers and speed decay factors. To simulate the system’s effective dynamics, we focused exclusively on high-adhesion B cell clusters which are modeled as circles with radius *r* and size *πr*^2^, treating the low-adhesion cells as a dense, unmodeled background. The self-organization process was modeled as a random walk of these B cell clusters within the background, where clusters merge upon contact to form a single cluster whose size equals the sum of the merged clusters. Each cluster performs a random walk with a speed of *s* = *s*_0_ · (*r/r*_0_)^−*x*^, where *s*_0_ and *r*_0_ are the speed and radius of a single cell, *r* is the cluster radius, and *x* determines how speed decays with cluster size.

Since the simulation plane is dense with either a high adhesion B-type cell, or a background low-adhesion cell, the size of the simulated plane is equal to 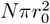 for *N* cells and initial cluster radius *r*_0_, with periodic boundary conditions.

The induced setup starts without high-adhesion cells, introducing a single high-adhesion cell every *k* time steps at a location chosen uniformly at random within the simulation plane. Conversely, the constitutive setup initializes all high-adhesion cells at time zero (at random locations), effectively making it the induced setup with *k* = 0.

Our simulations use the following default parameters: number of cells *N* = 1000, fraction of core cells = 0.3, k = 15, x = 3, simulation time steps = 22000, single cell radius *r*_0_ = 10, single cell speed *s*_0_ = 100, number of instantiations = 100.

### 4.2 Mammalian cell culture

Mouse L929 fibroblast cell lines used in the study were engineered as previously described [23]. Engineered cell lines were cultured in DMEM (Thermo Fisher Scientific 10566016) supplemented with 10% fetal bovine serum (UCSF Cell Culture Facility) in T25 flasks. Prior to kinetic imaging, each cell line was dissociated using 1x TrypLE (Thermo Fisher Scientific 12605028). Engineered cells were seeded in low attachment round bottom plates (Nexcelom ULA-384U-010) at either 300 total cells or 900 total cells with varying ratios of A-type and B-type cells (1:9, 3:7, 1:1, 7:3, 9:1). After seeding, the cells were centrifuged at 100 *× g* for 10 seconds prior to imaging. Each condition was seeded in quadruplicate.

### 4.3 Automated confocal microscopy and image analysis

Kinetic imaging was conducted using an Opera Phenix automated confocal microscope (PerkinElmer) using Harmony 4.9 software (PerkinElmer). Image stacks spanning a z-distance of 120 µm were acquired at one hour intervals over a period of 21 hours for each well of the 384 well plate. Subsequently, each experiment was analyzed using image analysis pipelines generated using Harmony 4.9 image analysis software using 3D analysis.

In the synNotch induced circuit, IFP positive receiver cells expressing anti-CD19 synNotch were identified using the ‘Find Nuclei’ building block. BFP/CD19 positive sender cells were identified using the same method as IFP positive receiver cells, but on the BFP channel. Subsequently, the mask for both IFP and BFP positive cells was reduced by 12 px to ensure no overlap between the volumetric masks. GFP intensity was quantified from the IFP positive cells using the ‘calculate intensity properties’ building block and GFP positive cells were identified via intensity thresholding. Positional information and morphological properties of each cell in all populations were quantified using the ‘calculate position properties’ and ‘calculate morphology properties’ building blocks. All data was output to.csv files for further analysis. For experiments to assess constitutive expression and sorting of L929s expressing high levels of E-cadherin, IFP positive cells and BFP positive cells were identified using the ‘Find Nuclei’ building block. Positional information and morphological properties of each cell in all populations were quantified using the ‘calculate position properties’ and ‘calculate morphology properties’ building blocks. All data was output to.csv files for further analysis.

### 4.4 Identification and quantification of cores

For the experimental microscopy images, at every time point we construct a graph *G* where the nodes represent high adhesion cells and are located at the coordinates of the corresponding cells. An edge between two nodes *B*_1_, *B*_2_ exists if they are in direct contact with each other, approximated as: dist(*B*_1_, *B*_2_) *≤* (1 + *ϵ*) ·(radius(*B*_1_) + radius(*B*_2_)), where dist() is the Euclidean distance, the radii are estimated as the cubic root of the measured cell volumes, and *ϵ* = 0.1 is a small constant that takes into account the uncertainty in the radii estimation. The number of cores is then computed as the number of connected components in *G*. To count cores in a manner robust to outlier cells (and avoid the counting of single cells as clusters), only clusters with a volume *v*_cluster_ *>* 3 · *v*_cell_ are considered cores, where *v*_cell_ is calculated as the median value of high adhesion B cell volumes at the initial time point across all trials and conditions.

In the computational models the core quantification method is similar to the experimental analysis: In the Potts model, we construct a regular graph where each grid point occupied by a high adhesion B cell is a node. Two nodes are connected if they are nearest neighbors on the grid, and the number of cores corresponds to the number of connected components in the graph. In the stochastic cluster model, cores are defined as clusters with at least 5% of the total B-type cell volume at a given time step, ensuring robustness against single high-adhesion cells introduced periodically.

### 4.5 Code Availability

The code used to generate all figures and results presented in this study is available at https://github.com/nitzanlab/coupling-signaling-adhesion. The repository includes scripts for running the Potts model, the stochastic cluster model, and the Morpheus simulations, as well as code for processing and analyzing experimental data. Detailed instructions for reproducing the simulations and analyses, along with dependencies and configuration files, are provided in the repository.

## Supplementary Information

**Supplementary Figure S1:**
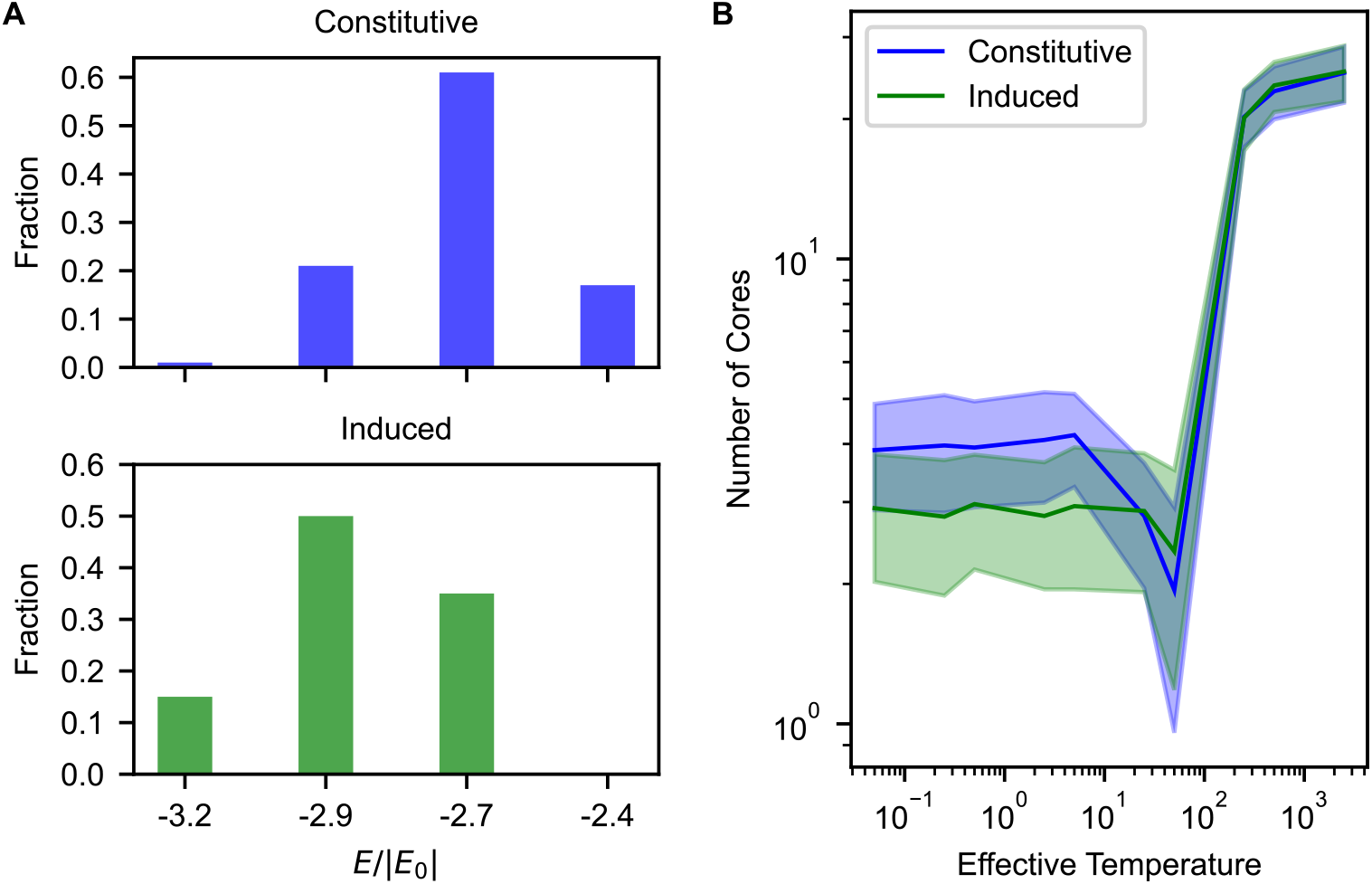
Simulations of the Potts model show that the induced system converges to a lower energy state than the constitutive system and remains robust across various effective temperatures. **(A)** Histogram of the configuration energy for the two circuits shows that the induced system tends to converge onto lower associated energy states relative to the constitutive system. The energy *E* is normalized by *E*0, the absolute value of the average initial energy of the constitutive system (random cellular configuration), calculated over 100 initializations. **(B)** Number of cores for the constitutive (blue) and induced (green) systems as a function of effective temperature (noise) (line: mean, shaded area: std). As the effective temperature of the simulated self-organization is varied, both the induced and constitutive systems exhibit a non-monotonic behavior where a minimal number of cores emerges at intermediate temperature values; while induced differentiation leads to reduced number of cores, low temperatures increase the probability that the systems will get kinetically trapped in suboptimal multi-core configurations. On the other hand, the behavior of the two systems in the high-temperature regime coincides as the high temperature disrupts any local structural convergence. Results are shown for 100 instantiations, Default parameters: number of cells = 200, fraction of core cells = 0.25, *T* = 5, binding energy: A-A, A-B = 10, B-B = 100, Monte Carlo simulation time steps = 400.

**Supplementary Figure S2:**
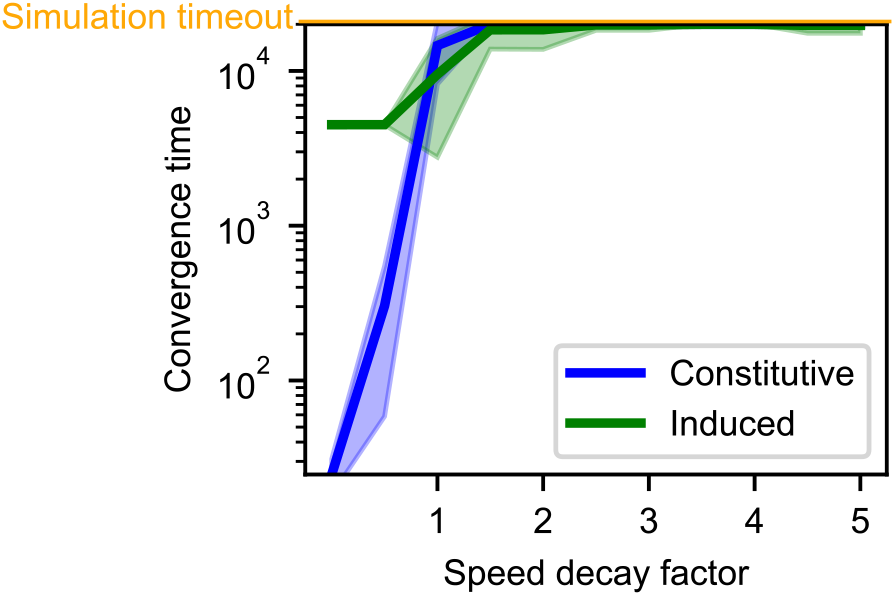
Convergence time increases with the speed decay factor *x* in the stochastic cluster model. The convergence time to a single core increases with the speed decay factor *x*, where the cluster speed *s* as a function of its radius *r* is modeled as *s ∝ r*^−*x*^ (line: mean, shaded area: std). For *x <* 1, the induced system takes longer to converge to a single core compared to the constitutive system, since it requires at least *NB* · *k* time steps for convergence, where *NB* is the number of B cells and *k* is the B cell activation rate. In the range 1 *< x <* 2, the induced system converges more quickly. Beyond *x >* 2, the constitutive system fails to converge to a single core in any instantiation, whereas the induced system achieves single-core convergence in a few cases. This result is further supported by Figure 2D, which shows that the induced system consistently achieves a lower number of cores. Default parameters: total cell number = 1000, fraction of core cells = 0.3, *k* = 15, number of instantiations = 100. See Section 4.1.2 for implementation details.

**Supplementary Figure S3:**
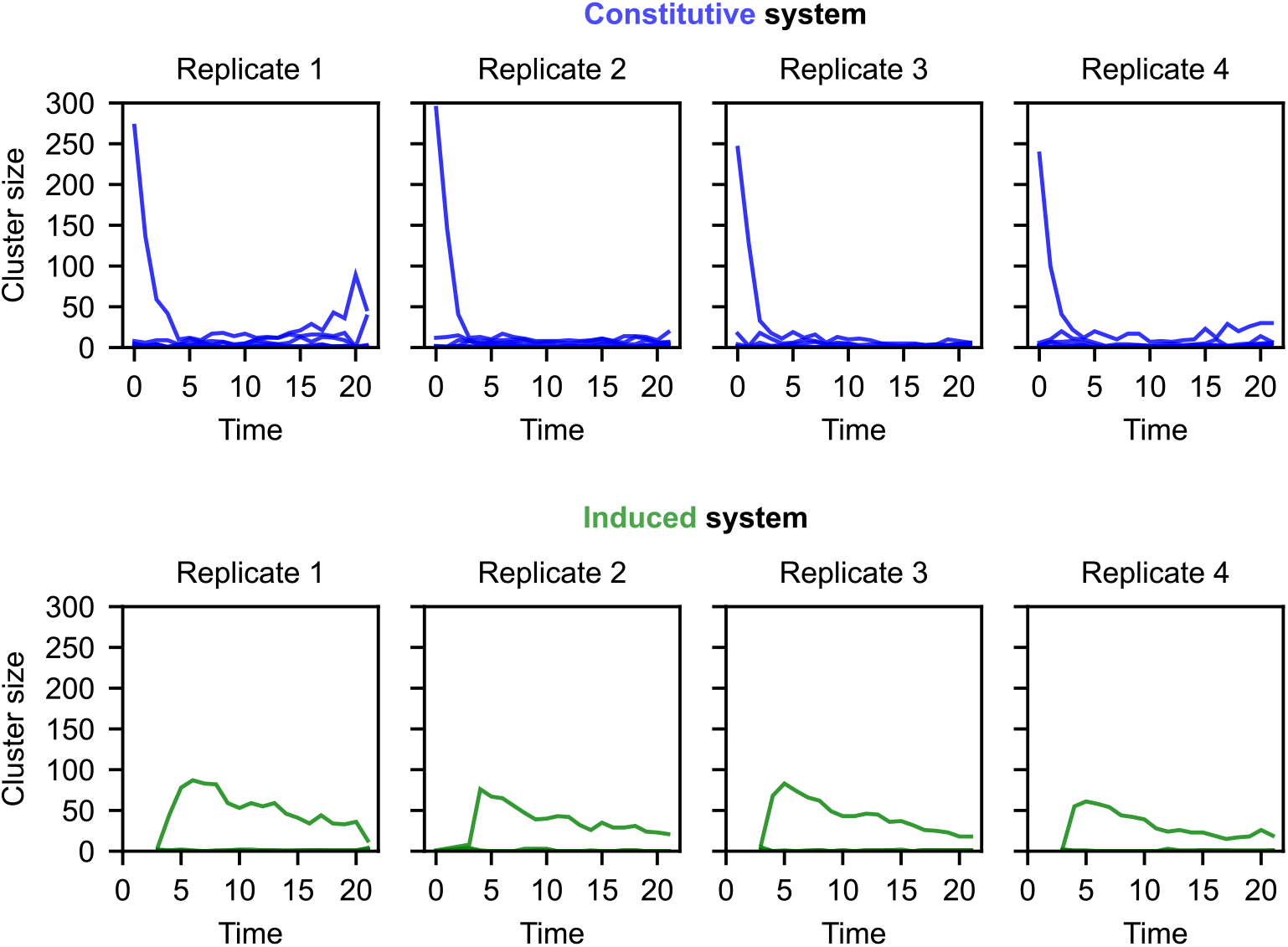
Cluster size dynamics in experimental observations show that the induced system converges to a single large core, whereas the constitutive system converges to multiple smaller cores. Cluster size over time for constitutive (A) and induced (B) systems. Different columns correspond to four replicates of the 900 cells, 1:1 cell ratio experiments. At each time point, clusters are identified according to the method described in Section 4.4. Each point on the graph represents the number of cells in a cluster at a specific time point. The lines connect cluster size points in an order-preserving manner (in terms of cluster size), representing the estimated trajectories of cluster sizes over time.

**Supplementary Figure S4:**
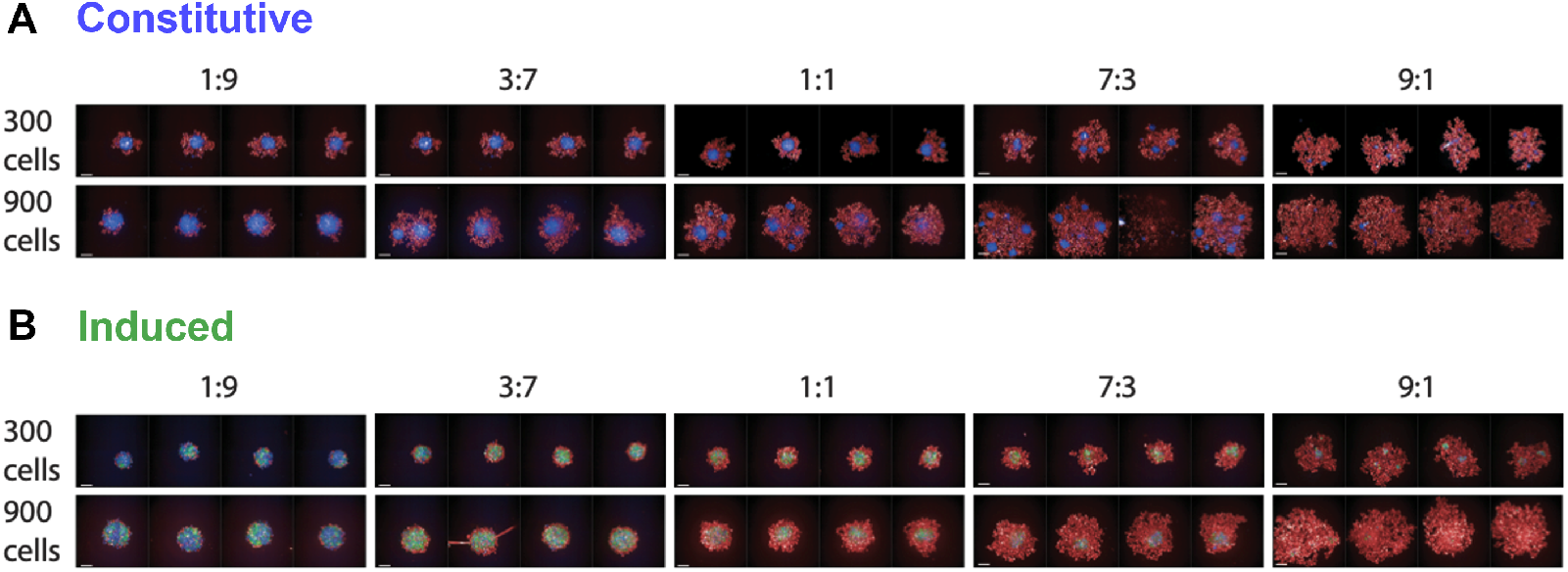
Experimental studies with synthetic core-shell systems show more stable convergence to a single core configuration of induced-adhesion compared to constitutive systems. (A,B) Final time experimental snapshot for constitutive (A) and induced (B) circuits, with different initial cell numbers (300 and 900), and different cell ratios (non-core to core cells: 1:9, 3:7, 1:1, 7:3, 9:1). Four replicates are shown for each experiment.

**Supplementary Figure S5:**
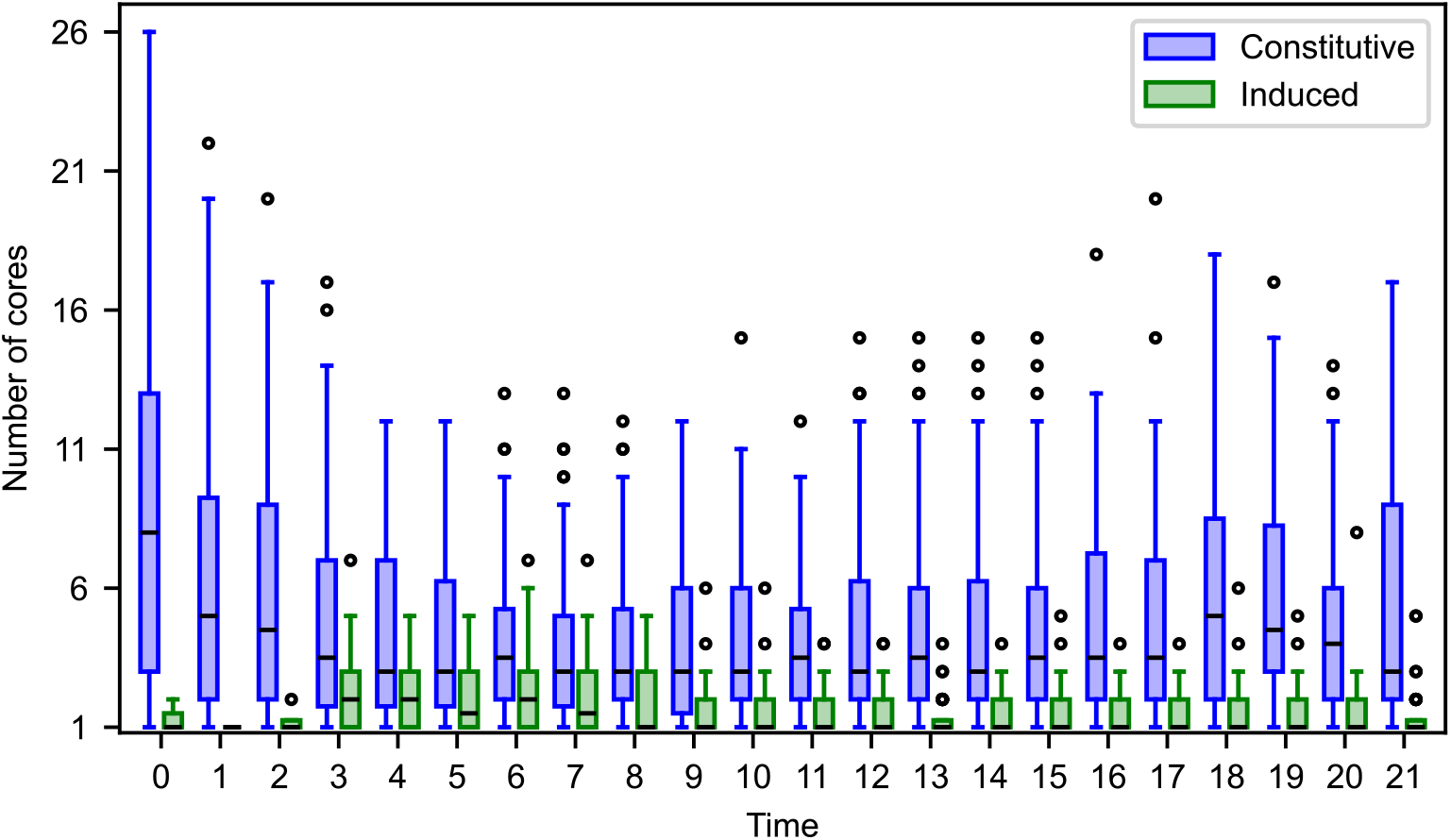
Experimental analysis of core counts across time points reveals that the induced system converges to fewer cores with lower variance compared to the constitutive system. Shown is a boxplot of the number of cores over all replicates and core fractions, as a function of time for the constitutive (blue) and induced (green) systems. The boxes extend from the lower to upper quartile values of the data, with a line at the median, whiskers extend from *Q*3 + 1.5 · (*Q*3 − *Q*1) to *Q*1 − 1.5 · (*Q*3 − *Q*1), where *Q*1 and *Q*3 are the first and third quartiles, and circles represent outlier points.

**Supplementary Figure S6:**
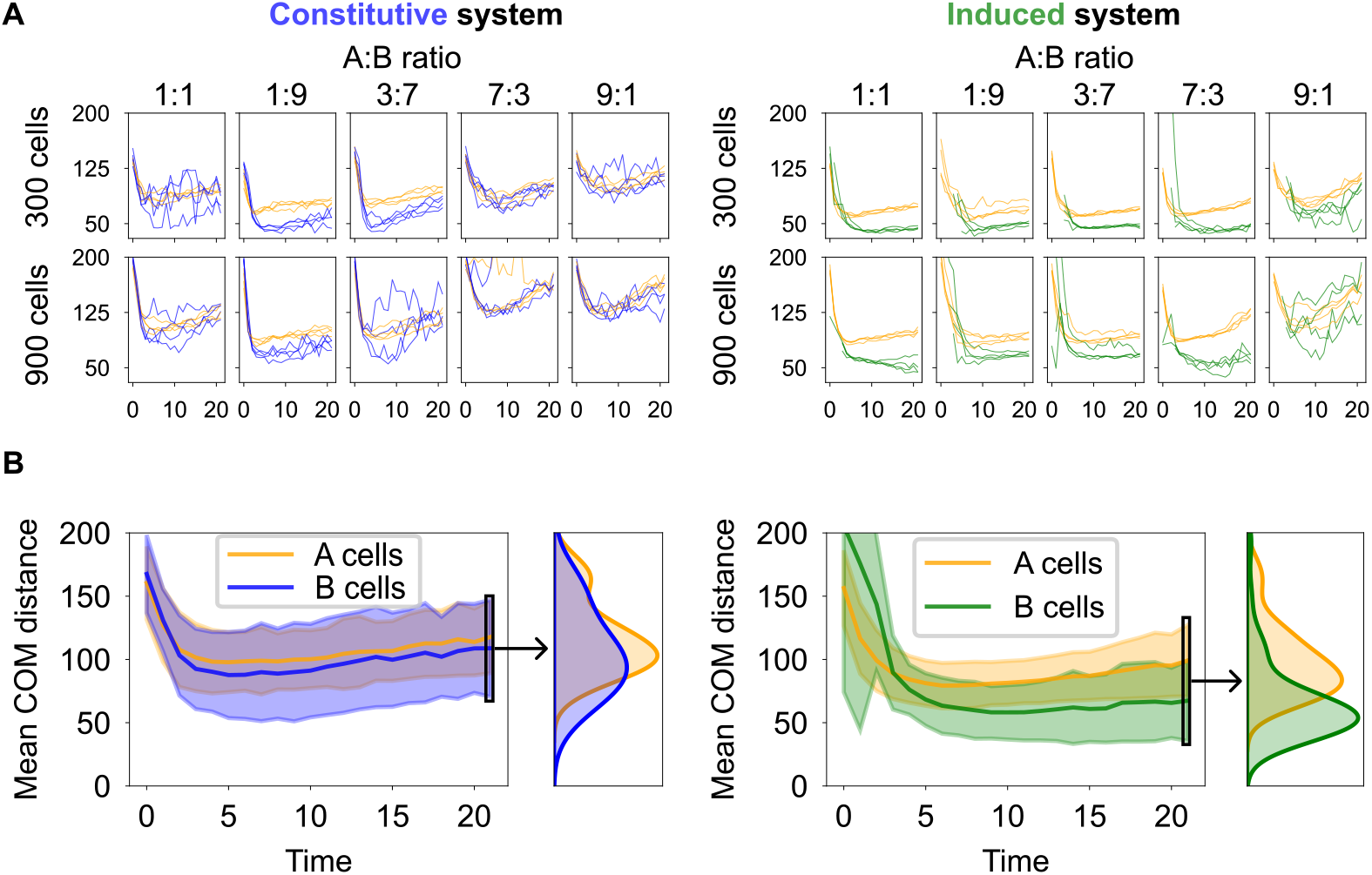
Experimental analysis of center-of-mass distance suggests that the induced system forms a compact core-shell structure, while core cells in the constitutive system remain more dispersed throughout the structure. **(A)** Average distance of low-adhesion A cells (yellow) and high-adhesion B cells (blue/green) from the spheroid’s center-of-mass (COM) over time for the constitutive (left) and induced (right) systems. Different rows (columns) represent varying total cell numbers (cell type ratios), and each trajectory corresponds to a single trial. In the engineered induced (constitutive) system, the structure forms and stabilizes after approximately 5.8 *±* 3.3 (1.9 *±* 0.8) hours, as defined by the earliest time point at which the mean distance from the COM is within 10% of the final value after 21 hours. **(B)** Mean COM distances averaged across all trials and conditions (shaded area: std). At the final time point, the mean COM distance of high-adhesion B cells in the induced system (67.4 *±* 31.7 µm) is lower compared to that in the constitutive system (108.8 *±* 38.4 µm). (Insets) In the induced system, the final mean distance from the center of mass (COM) reveals a clear separation between A and B-type cells, with B-type cells clustering more tightly to form a well-defined core-shell structure. In contrast, the constitutive system shows overlapping distributions of A and B-type cells, indicating weaker spatial segregation and less distinct core-shell organization.

**Supplementary Figure S7:**
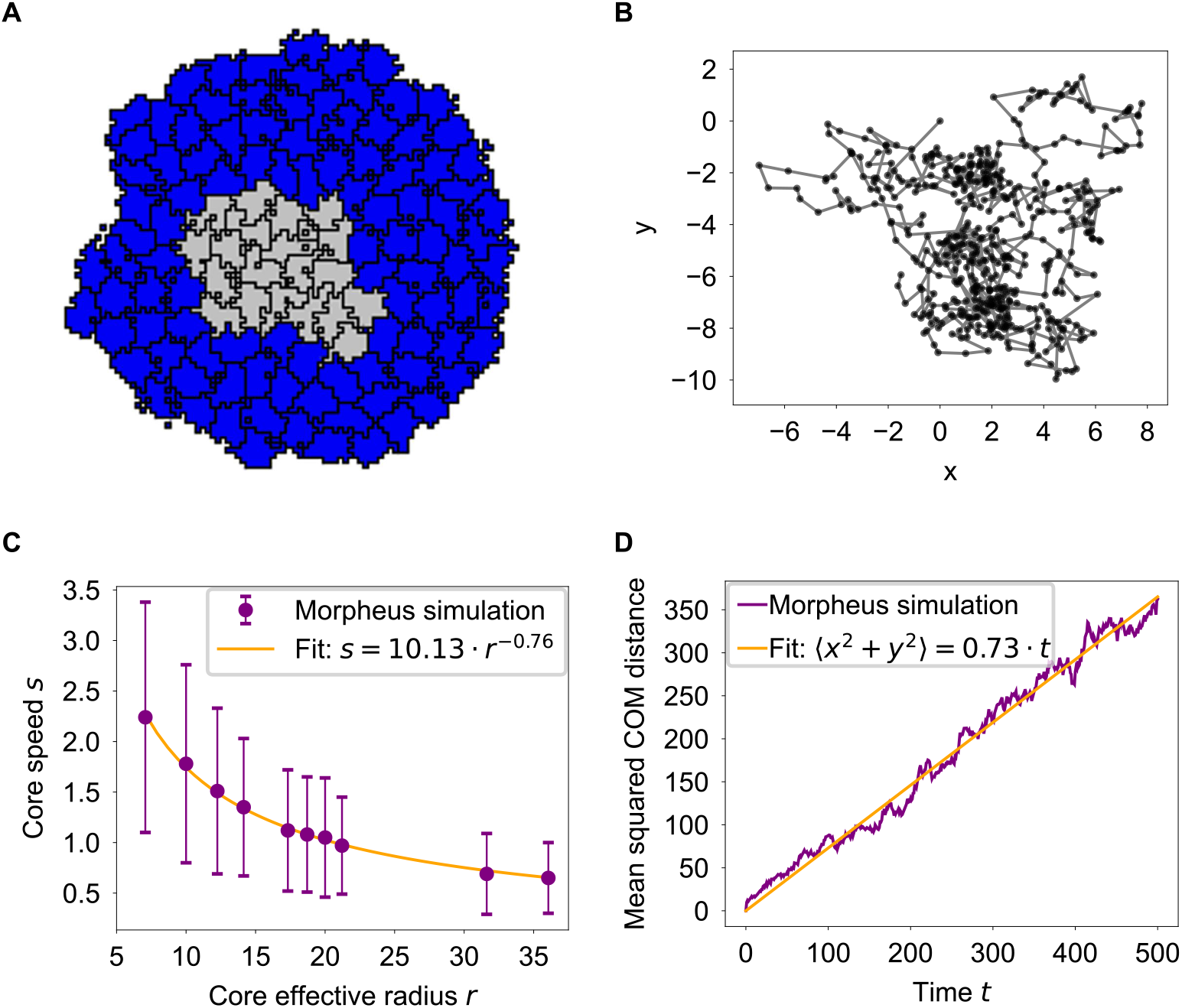
Simulations of core-shell dynamics in the Morpheus framework reveal a random walk pattern of the core, with core speed *s* decaying with its effective radius *r* as a power law *s ∝ r*^−*x*^. **(A)** Snapshot of the core-shell structure, where high adhesion cells (gray) form a core surrounded by low-adhesion cells (blue). **(B)** Trajectory of the core’s center of mass (COM), following a random walk. **(C)** The core’s COM speed plotted as a function of its effective radius, defined as the square root of the total area of high adhesion cells (points: mean, whiskers: std, line: the power-law fit). **(D)** The squared center of mass (COM) distance from the origin, averaged over 15 trials, increases linearly with the number of time steps, *< x*^2^ + *y*^2^ *>∝ t*, consistent with a random walk pattern. For details about the Morpheus framework see [27].

## References

[1] R. A. Foty and M. S. Steinberg. “The differential adhesion hypothesis: a direct evaluation”. In: Developmental Biology 278 (2005), pp. 255–263.

[2] D. Duguay, R. A. Foty, and M. S. Steinberg. “Cadherin-mediated cell adhesion and tissue segregation: qualitative and quantitative determinants”. In: Developmental Biology 253 (2003), pp. 309–323.

[3] A. Nose, A. Nagafuchi, and M. Takeichi. “Expressed recombinant cadherins mediate cell sorting in model systems”. In: Cell 54 (1988), pp. 993–1001.

[4] U. Tepass and K. P. Harris. “Adherens junctions in Drosophila retinal morphogenesis”. In: Trends in Cell Biology 17 (2007), pp. 26–35.

[5] A. Steinke et al. “Molecular composition of tight and adherens junctions in the rat olfactory epithelium and fila”. In: Histochemistry and Cell Biology 130 (2008), pp. 339–361.

[6] S. Katsunuma et al. “Synergistic action of nectins and cadherins generates the mosaic cellular pattern of the olfactory epithelium”. In: Journal of Cell Biology 212 (2016), pp. 561–575.

[7] H. Togashi et al. “Nectins establish a checkerboard-like cellular pattern in the auditory epithelium”. In: Science 333 (2011), pp. 1144–1147.

[8] J. E. Cooke, H. A. Kemp, and C. B. Moens. “EphA4 is required for cell adhesion and rhombomere-boundary formation in the zebrafish”. In: Current Biology 15 (2005), pp. 536–542.

[9] S. Ohgo et al. “Analysis of hoxa11 and hoxa13 expression during patternless limb regeneration in Xenopus”. In: Developmental Biology 338 (2010), pp. 148–157.

[10] E. Cachat et al. “2- and 3-dimensional synthetic large-scale de novo patterning by mammalian cells through phase separation”. In: Scientific Reports 6 (2016), p. 20664.

[11] J. Davies. “Using synthetic biology to explore principles of development”. In: Development 144 (2017), pp. 1146–1158.

[12] M. Pour et al. “Emergence and patterning dynamics of mouse-definitive endoderm”. In: iScience 25 (2022), p. 103556.

[13] A. Ferrer-Vaquer, M. Viotti, and A.-K. Hadjantonakis. “Transitions between epithelial and mesenchymal states and the morphogenesis of the early mouse embryo”. In: Cell Adhesion and Migration 4 (2010), pp. 447–457.

[14] M. Viotti, S. Nowotschin, and A.-K. Hadjantonakis. “SOX17 links gut endoderm morphogenesis and germ layer segregation”. In: Nature Cell Biology 16 (2014), pp. 1146–1156.

[15] R. Niu et al. “Magnetic handshake materials as a scale-invariant platform for programmed self-assembly”. In: Proceedings of the National Academy of Sciences of the United States of America 116 (2019), pp. 24402–24407.

[16] Z. Zeravcic, V. N. Manoharan, and M. P. Brenner. “Size limits of self-assembled colloidal structures made using specific interactions”. In: Proceedings of the National Academy of Sciences of the United States of America 111 (2014), pp. 15918–15923.

[17] P. W. K. Rothemund. “Folding DNA to create nanoscale shapes and patterns”. In: Nature 440 (2006), pp. 297–302.

[18] Wendell A. Lim. “The emerging era of cell engineering: Harnessing the modularity of cells to program complex biological function”. In: Science 378.6622 (2022), pp. 848–852.

[19] Kosuke Mizuno, Tsuyoshi Hirashima, and Satoshi Toda. “Robust Tissue Pattern Formation by Coupling Morphogen Signal and Cell Adhesion”. In: EMBO Reports 25.11 (Sept. 2024), pp. 4803–4826. ISSN: 1469-3178. doi: 10.1038/s44319-024-00261-z. (Visited on 04/02/2025).

[20] Mher Garibyan et al. “Engineering Programmable Material-to-Cell Pathways via Synthetic Notch Receptors to Spatially Control Differentiation in Multicellular Constructs”. In: Nature Communications 15.1 (July 2024), p. 5891. ISSN: 2041-1723. doi: 10.1038/s41467-024-50126-1. (Visited on 04/02/2025).

[21] Toshimichi Yamada et al. “Synthetic Organizer Cells Guide Development via Spatial and Biochemical Instructions”. In: Cell 188.3 (Feb. 2025), 778–795.e18. ISSN: 00928674. doi: 10.1016/j.cell.2024.11.017. (Visited on 04/02/2025).

[22] Adam J. Stevens et al. “Programming Multicellular Assembly with Synthetic Cell Adhesion Molecules”. In: Nature 614.7946 (Feb. 2023), pp. 144–152. ISSN: 1476-4687. doi: 10.1038/s41586-022-05622-z. (Visited on 04/02/2025).

[23] S. Toda et al. “Programming self-organizing multicellular structures with synthetic cell-cell signaling”. In: Science 361 (2018), pp. 156–162.

[24] D. J. Cohen. “Sorting Things Out: Cell Sorting during Hydra Regeneration”. In: Biophysical Journal 113 (2017), pp. 2577–2578.

[25] G. Serrano Nájera and C. J. Weijer. “Cellular processes driving gastrulation in the avian embryo”. In: Mechanisms of Development 163 (2020), p. 103624.

[26] S. Bonner-Weir, B. A. Sullivan, and G. C. Weir. “Human Islet Morphology Revisited: Human and Rodent Islets Are Not So Different After All”. In: Journal of Histochemistry and Cytochemistry 63 (2015), pp. 604–612.

[27] Jörn Starruß et al. “Morpheus: a user-friendly modeling environment for multiscale and multicellular systems biology”. In: Bioinformatics 30.9 (Jan. 2014), pp. 1331–1332. ISSN: 1367-4803. doi: 10.1093/bioinformatics/btt772. eprint: https://academic.oup.com/bioinformatics/article-pdf/30/9/1331/48922498/bioinformatics\_30\_9\_1331.pdf. URL: https://doi.org/10.1093/bioinformatics/btt772.

[28] L. Morsut et al. “Engineering Customized Cell Sensing and Response Behaviors Using Synthetic Notch Receptors”. In: Cell 164 (2016), pp. 780–791.

[29] J. Tordoff et al. “Incomplete Cell Sorting Creates Engineerable Structures with Long-Term Stability”. In: Cell Reports Physical Science 2 (2021), p. 100305.

[30] M. D. Petkova et al. “Optimal Decoding of Cellular Identities in a Genetic Network”. In: Cell 176 (2019), 844–855.e15.

[31] J. O. Dubuis et al. “Positional information, in bits”. In: Proceedings of the National Academy of Sciences of the United States of America 110 (2013), pp. 16301–16308.

[32] T. Gregor et al. “Probing the limits to positional information”. In: Cell 130 (2007), pp. 153–164.

[33] L. Wolpert. “Positional information and the spatial pattern of cellular differentiation”. In: Journal of Theoretical Biology 25 (1969), pp. 1–47.

[34] Coralie Trentesaux et al. “Harnessing Synthetic Biology to Engineer Organoids and Tissues”. In: Cell Stem Cell 30.1 (Jan. 2023), pp. 10–19. ISSN: 1934-5909, 1875-9777. doi: 10.1016/j.stem.2022.12.013. (Visited on 04/03/2025).

[35] F. Graner and J. A. Glazier. “Simulation of biological cell sorting using a two-dimensional extended Potts model”. In: Physical Review Letters 69 (1992), pp. 2013–2016.

[36] J. A. Glazier, A. Balter, and N. J. Poplawski. “Magnetization to Morphogenesis: A Brief History of the Glazier-Graner-Hogeweg Model”. In: Mathematics and Biosciences in Interaction. Springer, 2007, pp. 79–106.

[37] Y. Zhang et al. “Computer simulations of cell sorting due to differential adhesion”. In: PLoS One 6 (2011), e24999.

[38] R. M. H. Merks and J. A. Glazier. “A cell-centered approach to developmental biology”. In: Physica A: Statistical Mechanics and its Applications 352 (2005), pp. 113–130.

[39] N. Mulberry and L. Edelstein-Keshet. “Self-organized multicellular structures from simple cell signaling: a computational model”. In: Physical Biology 17 (2020), p. 066003.

